# Bayesian joint species distribution model selection for community-level prediction

**DOI:** 10.1101/2022.05.03.490480

**Authors:** Malcolm Itter, Elina Kaarlejärvi, Anna-Liisa Laine, Leena Hamberg, Tiina Tonteri, Jarno Vanhatalo

## Abstract

Joint species distribution models (JSDMs) are an important conservation tool for predicting ecosystem diversity and function under global change. The growing complexity of modern JSDMs necessitates careful model selection tailored to the challenges of community prediction under novel conditions (i.e., transferable models). Common approaches to evaluate the performance of JSDMs for community-level prediction are based on individual species predictions that do not account for the species correlation structures inherent in JSDMs. Here, we formalize a Bayesian model selection approach that accounts for species correlation structures and apply it to compare the community-level predictive performance of alternative JSDMs across broad environmental gradients emulating transferable applications. We connect the evaluation of JSDM predictions to Bayesian model selection theory under which the log score is the preferred performance measure for probabilistic prediction. We define the joint log score for community-level prediction and distinguish it from more commonly applied JSDM evaluation metrics. We then apply this community log score to evaluate predictions of 1,918 out-of-sample boreal forest understory communities spanning 39 species generated using a novel JSDM framework that supports alternative species correlation structures: independent, compositional dependence, and residual dependence. The best performing JSDM included all observed environmental variables and multinomial species correlations reflecting compositional dependence within modeled community data. The addition of flexible residual species correlations improved model predictions only within JSDMs applying a reduced set of environmental variables highlighting potential confounding between unobserved environmental conditions and residual species dependence. The best performing JSDM was consistent across successional and bio-climatic gradients regardless of whether interest was in species- or community-level prediction. Our study demonstrates the utility of the community log score to quantify differences in the predictive performance of complex JSDMs and highlights the importance of accounting for species dependence when interest is in community composition under novel conditions.

## Introduction

Joint species distribution models (JSDMs) are an important tool to predict community composition in an era of global change (Blowes et al., 2019). These predictions may be used to advance understanding of biodiversity change (Antão et al., 2022), identify at risk populations or natural communities (Araújo et al., 2011), and inform conservation management decisions (Guisan et al., 2013; Pollock et al., 2020). The growing complexity of contemporary JSDMs, which allow for the inclusion of correlation among species, hierarchical estimation of species-specific environmental responses informed by functional traits and phylogeny, as well as spatio-temporal correlation among sample units (Pollock et al., 2014; Warton et al., 2015; Ovaskainen et al., 2017; Clark et al., 2017), combined with the challenges of making predictions under novel conditions, necessitates careful model selection to ensure meaningful predictions of communities under global change (Roberts et al., 2017). JSDMs differ from traditional species distribution models (SDMs) in that they model the joint occurrence or abundance of species as a function of fixed environmental effects and residual species correlations that account for statistical dependence among species after controlling for the observed environment (Clark et al., 2014; Pollock et al., 2014; Warton et al., 2015).

As highlighted in recent work (Wilkinson et al., 2021), JSDMs allow for several types of prediction including individual species (marginal), all species simultaneously (joint), and simultaneous prediction of a subset of species conditional on a second, disjoint subset (conditional). JSDMs are expected to provide the greatest improvement in prediction (relative to SDMs) under the joint and conditional cases given that they account for the statistical dependence among species (Poggiato et al., 2021). This improvement is illustrated by an early JSDM study wherein summing the predicted abundance of tree species among independent SDMs (i.e., stacked SDMs) led to unrealistically high total abundance values relative to a JSDM which accounted for dependence among species induced by fixed growing space (Clark et al., 2014). In contrast, SDMs and JSDMs are expected to yield similar, or under certain model specifications, identical marginal predictions of species occurrence or abundance (Wilkinson et al., 2019, 2021; Poggiato et al., 2021). Despite this expectation, the predictive performance of JSDMs is often evaluated based on the marginal prediction of species even in cases where interest is in community composition (Caradima et al., 2019; Zurell et al., 2020).

Recent work highlights the diversity of JSDM prediction types along with statistical methods to evaluate their accuracy (Broms et al., 2016; Wilkinson et al., 2021). In general, measures of predictive accuracy should follow directly from the application of the model and the consequences (i.e., statistical loss) of making bad predictions (Gelman et al., 2014; Hooten and Hobbs, 2015). In the context of JSDMs, this means that if interest is in community composition, models should be evaluated based on joint or conditional predictions that depend on the presence/abundance of other co-occurring species (Wilkinson et al., 2021; Poggiato et al., 2021). Here, we formalize and demonstrate an approach for evaluating community-level predictions applying Bayesian JSDMs (i.e., joint and conditional joint prediction types *sec* Wilkinson et al., 2021). We focus on Bayesian JSDMs given both their prevalence (Hui, 2016; Ovaskainen et al., 2017; Clark et al., 2017) and their capacity to accommodate and quantify varying sources of uncertainty in data and models (Cressie et al., 2009).

Statistical measures (i.e., scoring statistics) to evaluate community-level predictions include community dissimilarity indices (e.g., Bray-Curtis dissimilarity, Jaccard distance) and likelihood-based methods including the joint log score (Wilkinson et al., 2021). Community dissimilarity indices are well known in ecology and have been widely applied to assess the community-level predictive performance of JSDMs or stacked SDMs (D’Amen et al., 2015; Broms et al., 2016; Maguire et al., 2016; Norberg et al., 2019; Wilkinson et al., 2021). The joint log score is less well known in ecology and has been applied to assess the predictive performance of JSDMs in only a small number of cases (Harris, 2015; Ingram et al., 2020; Wilkinson et al., 2021). Despite its limited application to JSDMs, the log score is widely applied in Bayesian model selection when the objective is probabilistic prediction of out-of-sample data (Hooten and Hobbs, 2015). Probabilistic prediction refers to the case where modelers seek to make inference on the complete probability distribution of future data reflecting all sources of uncertainty, rather than a point estimate such as the mean (Vehtari and Ojanen, 2012). In this context, the log score is the preferred metric for Bayesian model selection given it is the unique local and proper scoring rule (Gelman et al., 2014; Hooten and Hobbs, 2015).

We define different ways to quantify the joint log score for community-level prediction including clarifying what is meant by an independent versus community log score (Wilkinson et al., 2021). We then apply the community log score to compare alternative JSDMs and identify the best performing model for community-level prediction of 1,918 out-of-sample boreal forest understory plant communities comprising 39 species. The applied model framework extends existing JSDM approaches (e.g., the Hierarchical Model of Species Communities; Ovaskainen et al., 2017) to account for compositional dependence commonly encountered in vegetation survey data in addition to residual correlation among species. Candidate models include nested sets of environmental variables and alternative species dependence structures. We further assess model transferability by evaluating the performance of alternative models in predicting individual species and communities across out-of-sample data spanning broad successional and bioclimatic gradients. In doing so, we demonstrate the flexibility and utility of the log score for Bayesian JSDM selection and highlight the role of species correlations in community-level predictions.

## Methods

### Log score calculation

The joint log score for community-level prediction has been defined in several existing JSDM studies (Harris, 2015; Ingram et al., 2020; Wilkinson et al., 2021). Here, we define different ways the joint log score may be constructed across species and sites in order to account for species and spatio-temporal dependence if they are included in the applied JSDM. We adapt definitions and notation from Hooten and Hobbs (2015) to facilitate connection to a comprehensive review of Bayesian model selection. The log score for general out-of-sample data is expressed as log[**y**^oos^|**y**^obs^] = log *∫*_Θ_ [**y**^oos^|***θ***_*k*_][***θ***_*k*_|**y**^obs^]*d****θ***_*k*_ where **y**^oos^ defines out-of-sample (oos) data, **y**^obs^ represents observed (obs) data, ***θ***_*k*_ represents parameters under a candidate model *k*, and [·] indicates a probability distribution (Hooten and Hobbs, 2015). We make predictions of out-of-sample data (a proxy for new data) conditional on our fitted model (as defined by the posterior distribution [***θ***_*k*_ | **y**^obs^]) and integrate over uncertainty in our model parameters reflected in the posterior distribution yielding the posterior predictive distribution. The joint log score for community prediction can be estimated in two ways for a single site. The first method is based on the posterior predictive distribution of the community: 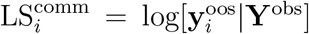. Here, 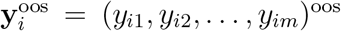 defines an out-of-sample local community consisting of *m* species (*i* = 1, …, *n*^oos^), while **Y**^obs^ is a matrix of observed natural communities (with dimension *n*^obs^ × *m*). In this case, 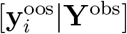 represents the joint distribution of species at site *i* including any species correlations included in the applied JSDM (e.g., **y**_*i*_ ∼ MVN(***µ***_*i*_, Σ) where Σ is an *m* × *m* covariance matrix). This is equivalent to the joint log score as defined in Wilkinson et al. (2021). The second method to estimate the joint log score treats species independently of one another: 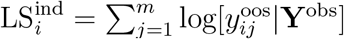 In this case, 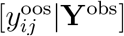 is the marginal distribution of species *j* at site *i* (e.g.,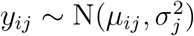) where 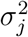 is the *j*th diagonal element of Σ). Note, the marginal distribution does not depend on other species as these have been integrated out. This is the so-called “independent” log score (Wilkinson et al., 2021). In the statistics literature, the latter is referred to as the point-wise log score (Gelman et al., 2014). Both versions of the joint log score can be calculated from a JSDM, but only the community log score includes modeled species correlations. When species are modeled independently of one another as in stacked SDMs, there is no difference between the two log scores because the joint distribution of species is equivalent to the product of their marginal distributions: 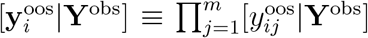 When interest is in community-level prediction applying a JSDM that includes statistical dependence among species, the community log score is preferred since it aligns with the modeler’s objective.

When multiple out-of-sample communities exist (as is likely the case), the joint log score (either the community or independent) may be summed across sites 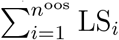 LS to estimatethe combined log score. This approach assumes that out-of-sample sites are statistically independent in time and or space. Additional approaches to construct the combined log score that take into account spatio-temporal correlation among sites if they are included in the applied JSDM are defined in the Supporting information (these are not applied in the current study given that we do not model spatio-temporal correlation among sites).

### Multinomial JSDM for compositional data

We define a multinomial data model within a common hierarchical Bayesian JSDM framework (Ovaskainen et al., 2017) to accommodate compositional community data frequently encountered in vegetative surveys wherein a fixed area plot is used to sample the count or percent cover of each species present. Similar versions of the multinomial model have been developed and applied to model microbiome data as well as plant compositional cover data, but have not been formally integrated within a JSDM framework (Xia et al., 2013; Damgaard, 2015, 2018). Under the multinomial model, a species’ relative abundance is estimated as a function of its expected success relative to the expected success of all species given local conditions. We apply the same notation for a local community as above with **y**_*i*_ denoting an *m*-dimensional vector of species’ relative abundances. The relative abundance of each species is modeled as,

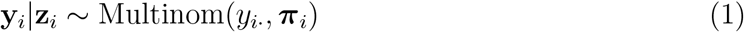

where **z**_*i*_ = (*z*_*i*1_, *z*_*i*2_, …, *z*_*im*_) defines an unobserved (latent) continuous variable, 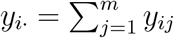 where *j* indexes species (*j* = 1, 2, …, *m*), and ***π***_*i*_ = (*π*_*i*1_, *π*_*i*2_, …, *π*_*im*_) where *π*_*ij*_ is the ex-pected probability of success of species *j* at site *i*. The expected probability of success is modeled applying a softmax function of **z**_*i*_,

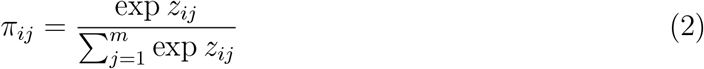

ensuring each *π*_*ij*_ ∈ (0, 1) and 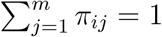. The *z*_*ij*_ variables are modeled as,

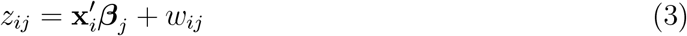

where **x**_*i*_ is a vector of site-level variables characterizing the local environment, ***β***_*j*_ is a vector of species-specific responses to each site-level variable, and *w*_*ij*_ is an additive species random effect accounting for variation not attributable to observed site conditions (reflecting random variation in species abundance or unobserved site conditions). Species-specific responses (***β***_*j*_ ‘s) are estimated applying a hierarchical prior allowing for partial pooling of data among species. A zero-centered Gaussian distribution is applied to model species random effects either independently or dependently (jointly) among species depending on the candidate model (see Model selection). Additional details on the JSDM structure including differences between independent and dependent random effect terms is provided in the Supporting information.

We implemented our model framework within the Hierarchical Model of Species Communities (HMSC) R package (Ovaskainen et al., 2017; Tikhonov et al., 2020, 2021). The HMSC package has a similar model structure to the one presented above, but does not include a multinomial data model (Eqtn. 1). Instead, we applied the well-established Poisson approximation to the multinomial to induce a multinomial likelihood for the linear predictor (**z**_*i*_), which uniquely defines species-specific probabilities (McCullagh and Nelder, 1989; Baker, 1994). Details on the Poisson approximation to the multinomial and its implementation within HMSC are provided in the Supporting information.

### Natural community data

We applied the multinomial JSDM to observations of Finnish boreal forest understory communities collected as part of national vegetation inventories. Understory vegetation was surveyed in 1985-1986, 1995, and 2006 across 1,700 unique plot locations established on mineral-soil in forested land. Plots were part of the systematic sampling network connected to the 8th Finnish National Forest Inventory (Reinikainen et al., 2000). This network consisted of clusters located 16 km apart in southern Finland, and 24 and 32 km apart in northern Finland along east–west and north–south axes, respectively. Each cluster consisted of four linearly located sampling sites 400 m apart in southern Finland and three sampling sites 600 m apart in northern Finland. Data includes 1,494 sites measured in 1985-1986, 1,673 sites in 1995, and 435 sites in 2006 (3,602 sites total). The survey in 2006 was part of the BioSoil project carried out under the Forest Focus scheme, a subset of the pan-European International Co-operative Programme on Assessment and Monitoring of Air Pollution Effects on Forests plot network (Level I, Lorenz and Fischer, 2013). This survey included only one site per cluster (hence the smaller sample size), but the spatial extent of the inventory was comparable to the 1985 and 1995 surveys and spanned the entirety of Finland (Fig. S.1). The surveys included observations of a total of 380 understory plant species. To ensure that meaningful values of species-specific model parameters could be estimated, we focused only on those species which occurred in at least five percent of sites in each inventory year reducing the total number of species to 39 (a common practice in high-dimensional JSDM settings, see Clark et al., 2017).

In all three surveys, vascular plant species were identified and each species’ percent cover (0.1 to 100 percent) was visually estimated within four permanent 2 m^2^ quadrats located 5 m apart within each site. There was no requirement that the percent cover values of all species sum to 100 percent given that a quadrat may not be fully occupied or may have a total cover greater than 100 percent if plants over top one another (a common occurrence in understory communities). To facilitate modeling, percent cover values were treated as numeric counts (percent cover values were rounded up to the nearest integer) representing the relative abundance of each species with the total abundance given by the sum across all species. When modeling understory communities, we calculated the average percent cover across the four sampled quadrats before converting percentages to estimates of relative and total abundance per site.

A number of environmental variables characterizing local conditions were measured along with understory community composition. The basal area of overstory trees at each site was estimated using measurements of stem diameter at breast height (Tomppo et al., 2011) and provides information on past forest management given basal area is inversely related to past harvest intensity (Fig. S.2). Shrub cover at each site was quantified as the projected percent cover of shrubs and 0.5-1.5 m tall trees within a 9.8 m radius circular plot centered on the permanent vegetation survey site. Soil fertility at each site was determined in the field using six ordinal classes based on vegetation (Cajander, 1949; Tomppo et al., 2011). For the purposes of this study, we re-classified these into two groups representing “high” and “low” fertility. Finally, growing degree days were estimated as the average annual sum of daily mean temperatures exceeding +5^*◦*^C per site over the decade preceding the inventory year based on 10 km^2^-resolution interpolated daily temperature values modeled by the Finnish Meteorological Institute (Venäläinen et al., 2005).

### Model selection

We defined eight candidate models applying alternative environmental variables and species random effect structures in combination with two alternative data models (Table 1): the multinomial model defined in Eqtn. (1); and a log-normal Poisson model, which assumed species’ relative abundances are independent conditional on **z**_*i*_,

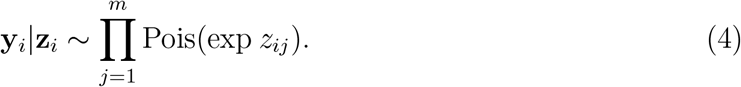

**Table 1:** Summary of candidate models. Variable abbreviations are defined as follows: BA = basal area per hectare, SoilFert = soil fertility, GDD = growing degree days, ShrubCover = percent shrub cover.

| Model | Environmental<br>variables ($\mathbf{x}_i$) | Species<br>dependence | Data<br>model |
| --- | --- | --- | --- |
| Stochastic-independent | None | Independent | Log-normal<br>Poisson |
| Management-independent | BA | Independent | Log-normal<br>Poisson |
| Environment-independent | BA, SoilFert,<br>GDD, ShrubCover | Independent | Log-normal<br>Poisson |
| Stochastic-compositional | None | Compositional | Multinomial |
| Management-compositional | BA | Compositional | Multinomial |
| Environment-compositional | BA, SoilFert,<br>GDD, ShrubCover | Compositional | Multinomial |
| Management-residual | BA | Compositional +<br>residual | Multinomial |
| Environment-residual | BA, SoilFert,<br>GDD, ShrubCover | Compositional +<br>residual | Multinomial |

Three environmental variable sets were tested. Stochastic models included an intercept term alone to estimate the mean abundance of each species across all sites. Management models included an intercept and basal area per hectare (BA) to account for time since forest overstory harvest. Finally, environment models included an intercept and BA along with growing degree days (GDD), soil fertility (SoilFert), and percent shrub cover (ShrubCover) to account for past management and local environmental conditions. A model selection approach identical to the one described herein was applied to identify the combination of management and environmental variables (including interactions) providing the best out-of-sample prediction. The final set of variables included the following terms: Intercept, SoilFert, BA, GDD, ShrubCover, SoilFert × BA, SoilFert × GDD, BA × GDD, SoilFert × BA × GDD.

Three species correlation structures were included across the eight candidate models. Independent models assumed no correlation among species and applied the log-normal Poisson data model (Eqtn. 4). Independent models represent multi-species distribution models under which species-specific responses to the environment are modeled hierarchically with no species correlation (Poggiato et al., 2021). (2) Compositional dependence models applied the multinomial data model (Eqtn. 1), which accounts for multinomial correlation among species. Note that multinomial correlation is constrained to be between -1.0 and 0.0 and accounts only for the fact that as the relative abundance of one species increases, the total relative abundance of all other species must decrease for a fixed community size (*π*_*ij*_ = 1 − Σ_*j′*≠*j*_ *π*_*ij′*_). We refer to the narrow form of species dependence captured by the multinomial data model as “compositional dependence.” (3) Residual dependence models applied the multinomial data model, but also allowed for residual correlation among species captured through a flexible dependent species random effect (supporting positive and negative pairwise correlations among species as opposed to the restrictive multinomial correlations). We refer to additional species dependence arising through the random effect term as “residual dependence” as it captures pairwise dependence among species after controlling for the effects of the environment. Model details are provided in the Supporting information.

We applied the community log score (as defined under Log score calculation) to assess the out-of-sample predictive performance of each candidate model. Specifically, we estimated the community log score for a subset of the 1985 and 1995 datasets including 1,918 out-of-sample natural communities by fitting each candidate model twice: once using the 2006 inventory data alone to predict 435 communities within the 1995 data (1995 | 2006), and once using the 1995 and 2006 inventory data combined to predict 1,483 communities within the 1985 data (1985 | 1995, 2006). The statistical basis for our out-of-sample predictive approach is provided in the Supporting information. To assess model transferability (i.e., predictive performance under novel conditions; Roberts et al., 2017), we compared the performance of a subset of four models (stochastic-independent, environment-independent, stochastic-compositional, environment-compositional) to predict individual species and communities across out-of-sample data representing different combinations of successional stage and climate. Specifically, we partitioned the 1985 and 1995 inventory data based on BA and regional bioclimatic conditions. The BA variable was divided into three ranges to represent management history: 0-10 m^2^/ha, 10-30 m^2^/ha, and *>* 30 m^2^/ha. The low BA range is indicative of understory communities early in ecological succession, while the high BA range is indicative of later successional communities given prolonged time since forest overstory harvest. We utilized three bioclimatic zones spanning the latitudinal gradient of Finland (south, mid, and north boreal) to represent regional climatic conditions (Table S.1, Fig. S.1; Ahti et al., 1968). This resulted in eight data partitions (the highest BA range was dropped for the north boreal zone given lack of sufficient out-of-sample data) with a minimum of 54 communities (mid-boreal, high basal area) and a maximum of 565 communities (south-boreal, mid basal area; Table S.2). Within each partition, we estimated the mean log score (across sites) under the four models for each species and the overall community.

## Results

Community log scores for each alternative model are provided in Fig. 1. Based on the log score results, the environment-compositional model including the full set of environmental predictors and compositional dependence among species had the highest joint predictive performance across the 1,918 out-of-sample communities. The top three performing models all included the complete set of environmental predictors (environment models). The next best three models included forest overstory density (BA) as the only environmental predictor (management models). The poorest performing models were those without any environmental predictors included (stochastic models). The top-performing model included compositional species dependence alone, while the second-best performing model did not include any species dependence. The environment-residual model with both compositional and residual species dependence had the poorest performance of the three models including the full set of environmental predictors. In-sample and out-of-sample posterior predictive model checks under the best predicting environment-compositional model revealed high correlation with observed data and species-level R-squared values exceeding 0.60 for the majority of species across all inventory years (Table S.4, Fig. S.3; see Supporting information for model checking details).

**Figure 1:**
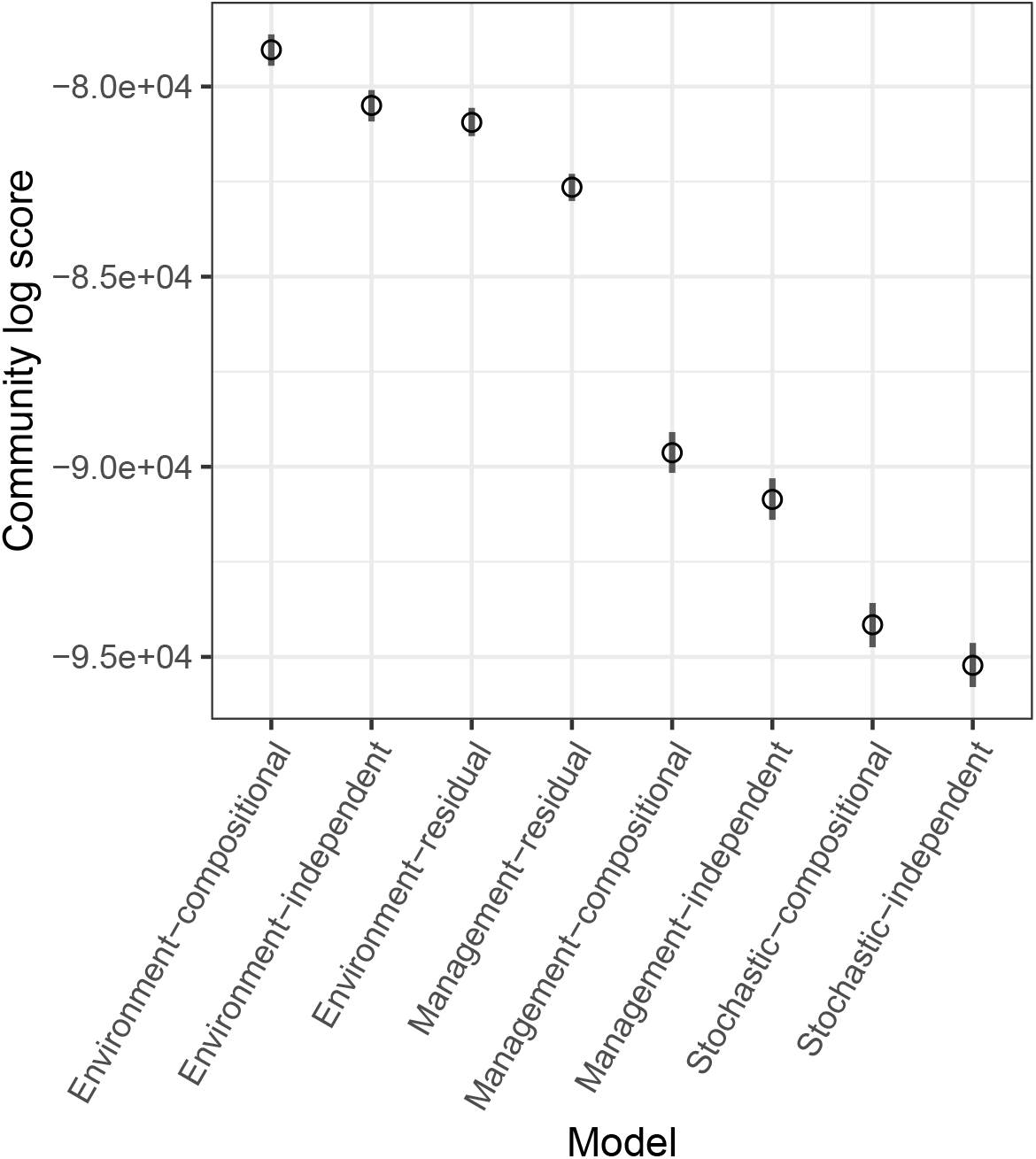
Community log scores for alternative models (as calculated by Eqtns. S.4 or S.5). Points represent bootstrapped median values, while lines indicate bootstrapped 95 percent confidence intervals. Models are ordered from highest to lowest log score with higher values indicating better out-of-sample prediction. Models names are defined in Table 1.

Similar to the community log score results for the complete out-of-sample data (Fig. 1), the environment-compositional model had the highest community-level predictive performance across all out-of-sample data partitions (Fig. 2). The differences between the predictive performance of the two environment models including the full set of environmental predictors and either compositional dependence (environment-compositional) or inde-pendence (environment-independent) among species were small with 95 percent confidence intervals for the two models overlapping in all partitions. There was clear separation of the two environment models relative to the two stochastic models (stochastic-independent, stochastic-compositional), but the 95 percent confidence intervals for all four models overlapped in half of the out-of-sample data partitions (Fig. 2). The environment-compositional model also had the highest predictive performance for the majority of species within every out-of-sample data partition. The environment-independent model was the best predicting model for the second-highest number of species in each partition followed by the stochastic-compositional model (Fig. 3).

**Figure 2:**
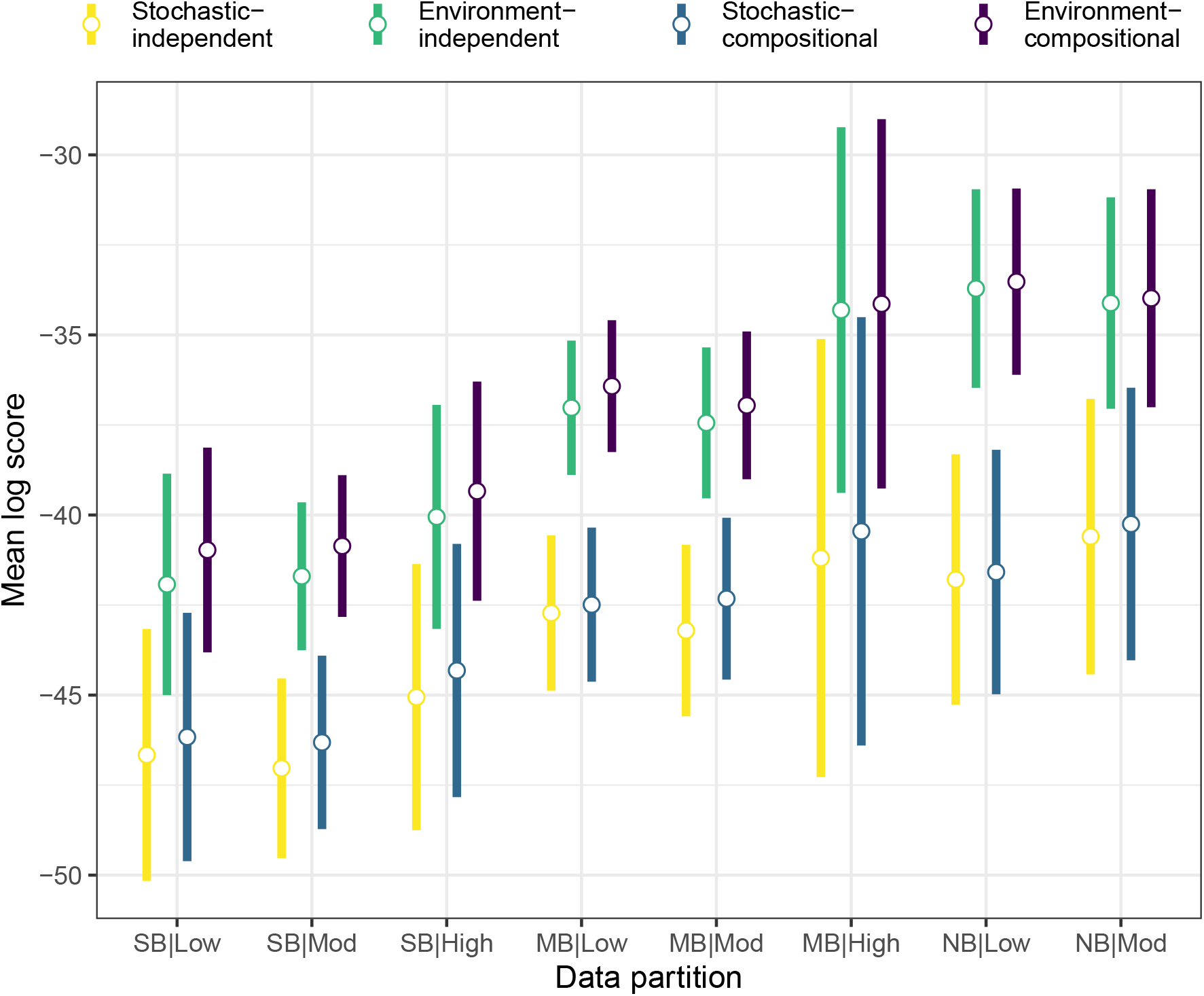
Mean, per-site community log scores for community-level prediction under the four alternative models applied within each out-of-sample data partition (as calculated by Eqtn. S.6). Points represent the mean across all sites within a partition, while lines indicate 95 percent confidence intervals. Data partition labels are defined as: bioclimatic zone | basal area (BA). Bioclimatic zones: SB = south boreal, MB = mid boreal, NB = north boreal;BA levels (m^2^·ha^*−*1^): Low = [0-10), Mod = [10-30), High = ⩾30.

**Figure 3:**
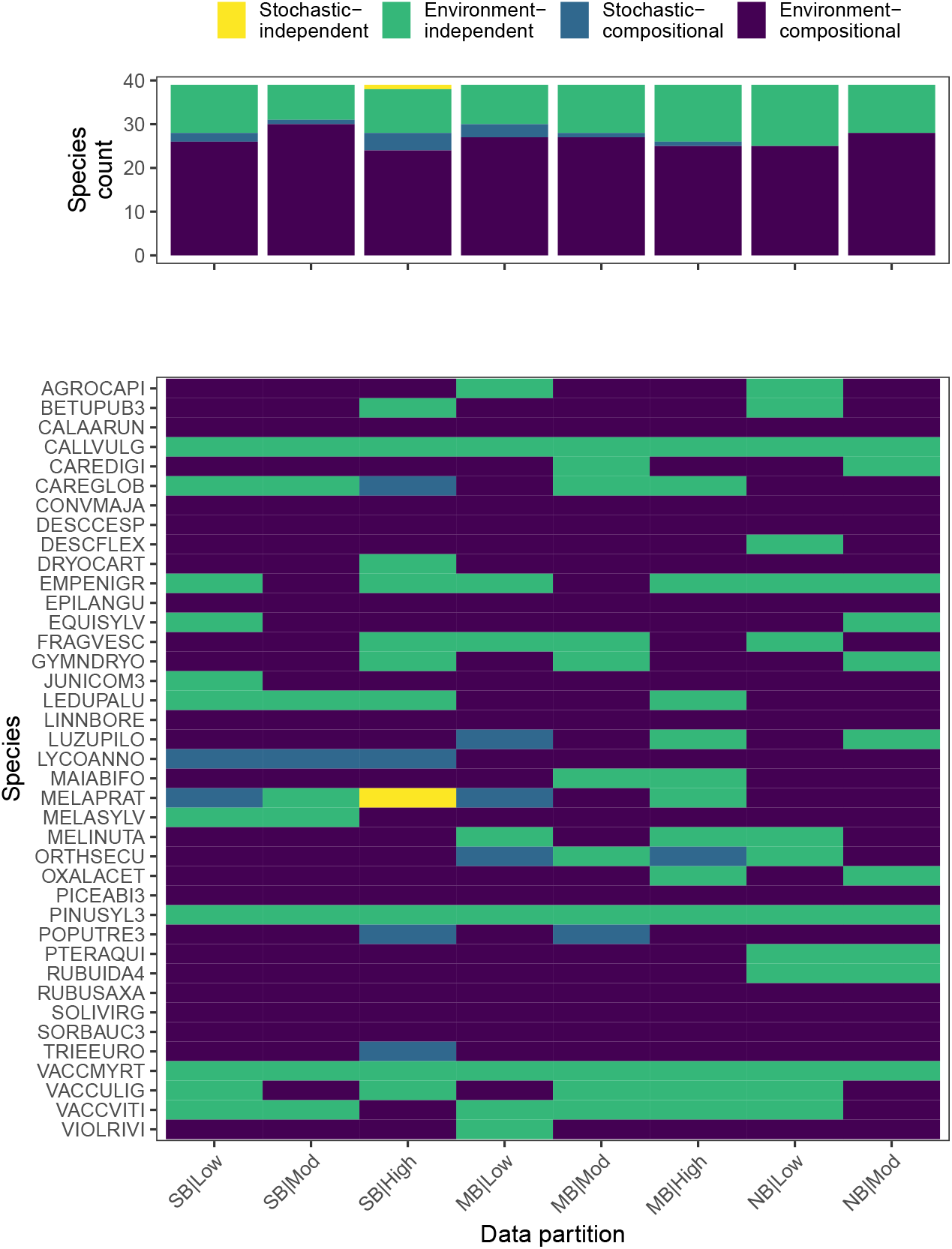
Summary of mean, per-site independent log scores for species-level prediction under the four alternative models applied within each out-of-sample data partition (as calculated by Eqtn. S.7) including: the number of species for which each model had the highest mean log score (upper panel) and the model with the highest mean log score for each species-by-partition combination (lower panel). Data partition labels are defined as: bioclimatic zone | basal area (BA). Bioclimatic zones: SB = south boreal, MB = mid boreal, NB = north· boreal; BA levels (m^2^ ha^*−*1^): Low = [0-10), Mod = [10-30), High = ⩾30. Species codes aredefined in Table S.3.

## Discussion

We found clear evidence for the inclusion of the complete set of environmental variables for the prediction of boreal forest understory communities (Fig. 1). However, the role of species dependence in community prediction was more nuanced. Species dependence was included in the alternative models in three ways: (1) no dependence (independent models), compositional dependence alone (compositional models), (3) compositional and residual dependence allowing for both positive and negative species correlations (residual models; Tab. 1). The model including both the full set of environmental predictors and compositional species dependence alone, had the best predictive performance as measured by its community log score (Fig. 1). Interestingly, the relative importance of alternative species dependence structures changed when forest density alone was used to represent the environmental filter (management models). Among the management models, there was statistical support for including both compositional and residual species dependence to predict community composition as evidenced by the ordering of the models: management-residual *>* management-compositional *>* management-independent (Fig. 1). This ordering is what we would expect to find based on ecological theory. That is, community composition related to complex combinations of abiotic and biotic filters, here potentially reflected by the compositional and residual correlations among species (Vellend, 2016). While the ordering of the simple management models matches ecological expectations, when compared to the ordering among the more complex environment models it hints at a broader and known limitation of JSDMs: the confounding of unobserved environmental effects and species dependence (Poggiato et al., 2021). Specifically, there was strong evidence that including residual species dependence improved community-level prediction when environmental variation was poorly captured as under the management models (Fig. 1). When the full set of environmental predictors was included, however, there was near equal predictive performance of independent and residual dependence models despite the former assuming no species dependence and the latter including both compositional and residual species dependence (Fig. 1). This suggests that the variability accounted for by residual species dependence was captured by the additional fixed environmental effects in the three environment models. This should not be interpreted as the lack of complex interactions among species in boreal forest understory communities; rather, it indicates that residual dependence among species in modeled communities was limited after controlling for the strong effects of the observed environment or it may be due to confounding of the flexible residual species random effects and observed environmental variation (Van Ee et al., 2022).

There are a number of reasons why including compositional dependence among species would improve community-level prediction in boreal forest understory communities as indicated by the model selection results. First, such dependence reflects the nature of the observations. Understory communities were measured using fixed area, 2 m^2^ quadrats, with high relative abundance of a given species resulting in less available growing space and other resources for other species in the community. This type of resource limitation is observed regularly in forest ecosystems and is the basis for common conceptual models of forest development driven by density-dependent competition (e.g., Reineke’s self-thinning law; Reineke, 1933). Although compositional dependence does not necessarily indicate resource competition among species, model predictions combined with previous experimental results suggest such a connection may exist in the modeled boreal forest understory communities (Gundale et al., 2012).

The full environment model with compositional species dependence also had the best performance in terms of predicting communities and individual species across the novel conditions reflected in out-of-sample data partitions spanning broad successional and bio-climatic gradients (Figs. 2, 3). The applied out-of-sample data partitions emulate block cross-validation strategies advocated for in recent work to assess the transferability of JS-DMs (Roberts et al., 2017). Based on recent work demonstrating spatio-temporal changes in the processes underlying community composition (Jabot et al., 2020), we expected alternative models to have varying predictive performance conditional on the successional stage and relative harshness of the regional climate. Contrary to our expectations, however, a consistent ordering of models emerged across all out-of-sample data partitions: environment-compositional *>* environment-independent *>* stochastic-compositinal *>* stochastic-independent (Fig. 2; note the relative differences among models varies by partition). The strong predictive performance of both the environment-compositional and environment-independent models across a range of successional stages and bioclimatic conditions at both the community and individual species scales highlights the robustness of model predictions when the full set of environmental variables is included and points to higher potential for model transferability. Note that the predictive performance for individual species is assessed based on marginal predictions (see Supporting information), providing evidence that the environment-compositional model is preferred for predicting both individual species and community composition.

The community log score (as defined under Log score calculation) provides a probabilistic, local, and proper metric to assess the predictive performance of JSDMs when interest is in community composition. The current study demonstrates the practical application of the community log score for community-level prediction and highlights its capacity to quantify differences in the predictive performance of highly structured JSDMs including complex species dependence structures. Further, the application of the log score to assess predictive performance across out-of-sample data partitions spanning broad successional and bioclimatic gradients demonstrates its utility for transferable model selection where the goal is to identify the best performing model under novel conditions. Although not the direct objective of the current analysis, the multinomial JSDM provides a multivariate approach for modeling compositional data commonly encountered in vegetation surveys. Given its implementation within the more general HMSC framework, the model can deal with a large number of species and locations and allows additional information on functional traits and phylogeny to be incorporated as predictors of environmental responses (Ovaskainen et al., 2017). As such, the model framework combined with the Bayesian model selection approach applied here, constitutes a step forward in our ability to predict plant communities under global change.

## Supporting information

Technical appendix

