## Supplemental tables and figures for "Bayesian joint species distribution model selection for community-level prediction"

Table S.1: Summary of climate within the three bioclimatic zones used to partition data in the decade preceding each understory inventory (1985, 1995, 2006). Mean values are presented for understory inventory sample sites with standard errors in parentheses. Growing degree days is defined as the 10-year moving average of the total number of days with a daily mean temperature greater than 5 °C during the preceding decade.

| Bioclimatic zone | Decade | Annual precipitation (mm) | Annual temperature (°C) | Growing degree days |
|---|---|---|---|---|
| South boreal | 1975-1984 | 747.4 (1.6) | 3.3 (0.03) | 1212.5 (2.4) |
| | 1985-1994 | 768.4 (1.8) | 3.5 (0.03) | 1215.5 (2.5) |
| | 1996-2005 | 781.8 (1.4) | 4.3 (0.03) | 1354.1 (2.6) |
| Mid boreal | 1975-1984 | 714.1 (2.4) | 1.6 (0.03) | 1020 (2.9) |
| | 1985-1994 | 746.6 (2.7) | 1.8 (0.04) | 1023.3 (2.8) |
| | 1996-2005 | 778.5 (2.6) | 2.6 (0.03) | 1155.3 (2.8) |
| North boreal | 1975-1984 | 641 (5.1) | -1.2 (0.05) | 721.5 (6.8) |
| | 1985-1994 | 664 (4.8) | -0.7 (0.05) | 746.3 (6.3) |
| | 1996-2005 | 719.7 (4.7) | -0.1 (0.05) | 873.1 (7.3) |

Table S.2: Number of out-of-sample natural communities in each data partition (BA = basal area) used to assess alternative model performance across successional and bioclimatic gradients

| Bioclimatic zone | Low BA [0-10) $m^2 \cdot ha^{-1}$ | Moderate BA [10-30) $m^2 \cdot ha^{-1}$) | High BA $\geqslant 30$ $m^2 \cdot ha^{-1}$ | Total |
|---|---|---|---|---|
| South boreal | 266 | 565 | 126 | 957 |
| Mid boreal | 295 | 370 | 54 | 719 |
| North boreal | 127 | 114 | NA | 241 |
| Total | 688 | 1049 | 180 | 1917 |

Table S.3: Species code definitions for all 39 study species

| Species code | Species |
|---|---|
| AGROCAPI | *Agrostis capillaris* |
| BETUPUB3 | *Betula pubescens* |
| CALAARUN | *Calamagrostis arundinacea* |
| CALLVULG | *Calluna vulgaris* |
| CAREDIGI | *Carex digitata* |
| CAREGLOB | *Carex globularis* |
| CONVMAJA | *Convallaria majalis* |
| DESCCESP | *Deschampsia cespitosa* |
| DESCFLEX | *Deschampsia flexuosa* |
| DRYOCART | *Dryopteris carthusiana* |
| EMPENIGR | *Empetrum nigrum* |
| EPILANGU | *Epilobium angustifolium* |
| EQUISYLV | *Equisetum sylvaticum* |
| FRAGVESC | *Fragaria vesca* |
| GYMNDRYO | *Gymnocarpium dryopteris* |
| JUNICOM3 | *Juniperus communis* |
| LEDUPALU | *Ledum palustre* |
| LINNBORE | *Linnaea borealis* |
| LUZUPILO | *Luzula pilosa* |
| LYCOANNO | *Lycopodium annotinum* |
| MAIABIFO | *Maianthemum bifolium* |
| MELAPRAT | *Melampyrum pratense* |
| MELASYLV | *Melampyrum sylvaticum* |
| MELINUTA | *Melica nutans* |
| ORTHSECU | *Orthilia secunda* |
| OXALACET | *Oxalis acetosella* |
| PICEABI3 | *Picea abies* |
| PINUSYL3 | *Pinus sylvestris* |
| POPUTRE3 | *Populus tremula* |
| PTERAQUI | *Pteridium aquilinum* |
| RUBUIDA4 | *Rubus idaeus* |
| RUBUSAXA | *Rubus saxatilis* |
| SOLIVIRG | *Solidago virgaurea* |
| SORBAUC3 | *Sorbus aucuparia* |
| TRIEEURO | *Lysimachia europaea* |
| VACCMYRT | *Vaccinium myrtillus* |
| VACCULIG | *Vaccinium uliginosum* |
| VACCVITI | *Vaccinium vitis-idaea* |
| VIOLRIVI | *Viola riviniana* |

Table S.4: Species-level Bayesian $R^2$ summary. Values correspond to posterior mean with 95 percent credible interval in parenthesis.

| Species | Out-of-sample | | In-sample | |
|---|---|---|---|---|
| code | $\mathbf{Y}^{85}\|\mathbf{Y}^{95},\mathbf{Y}^{06}$ | $\mathbf{Y}^{95}\|\mathbf{Y}^{06}$ | $\mathbf{Y}^{95},\mathbf{Y}^{06}$ | $\mathbf{Y}^{06}$ |
| AGROCAPI | 0.9 (0.74,0.99) | 0.65 (0.3,0.97) | 0.93 (0.81,1) | 0.83 (0.53,0.99) |
| BETUPUB3 | 0.18 (0.1,0.29) | 0.06 (0.03,0.11) | 0.18 (0.11,0.29) | 0.11 (0.06,0.2) |
| CALAARUN | 0.98 (0.94,1) | 0.9 (0.72,0.99) | 0.99 (0.97,1) | 0.97 (0.9,1) |
| CALLVULG | 1 (0.99,1) | 0.99 (0.96,1) | 1 (0.99,1) | 0.99 (0.95,1) |
| CAREDIGI | 0.25 (0.15,0.43) | 0.21 (0.08,0.58) | 0.34 (0.2,0.57) | 0.33 (0.13,0.73) |
| CAREGLOB | 0.85 (0.72,0.97) | 0.57 (0.32,0.9) | 0.86 (0.73,0.97) | 0.62 (0.36,0.93) |
| CONVMAJA | 0.77 (0.54,0.97) | 0.56 (0.24,0.94) | 0.85 (0.67,0.98) | 0.72 (0.4,0.98) |
| DESCCESP | 0.96 (0.86,1) | 0.6 (0.27,0.95) | 0.97 (0.9,1) | 0.79 (0.49,0.99) |
| DESCFLEX | 0.95 (0.91,0.98) | 0.92 (0.84,0.98) | 0.95 (0.92,0.98) | 0.93 (0.86,0.99) |
| DRYOCART | 0.86 (0.72,0.98) | 0.71 (0.44,0.96) | 0.9 (0.79,0.99) | 0.8 (0.57,0.98) |
| EMPENIGR | 1 (0.99,1) | 1 (0.99,1) | 1 (0.99,1) | 0.99 (0.97,1) |
| EPILANGU | 0.84 (0.74,0.95) | 0.66 (0.44,0.9) | 0.88 (0.79,0.96) | 0.84 (0.68,0.97) |
| EQUISYLV | 0.74 (0.55,0.94) | 0.2 (0.04,0.66) | 0.77 (0.59,0.96) | 0.28 (0.09,0.78) |
| FRAGVESC | 0.19 (0.07,0.4) | 0.19 (0.07,0.54) | 0.25 (0.11,0.49) | 0.32 (0.11,0.75) |
| GYMNDRYO | 0.97 (0.89,1) | 0.76 (0.48,0.98) | 0.98 (0.92,1) | 0.8 (0.54,0.99) |
| JUNICOM3 | 0.43 (0.1,0.75) | 0.35 (0.13,0.76) | 0.45 (0.12,0.77) | 0.4 (0.16,0.83) |
| LEDUPALU | 0.99 (0.97,1) | 0.98 (0.92,1) | 0.99 (0.96,1) | 0.96 (0.83,1) |
| LINNBORE | 0.52 (0.41,0.68) | 0.71 (0.48,0.93) | 0.52 (0.42,0.66) | 0.68 (0.47,0.91) |
| LUZUPILO | 0.12 (0.1,0.14) | 0.17 (0.13,0.25) | 0.13 (0.11,0.15) | 0.24 (0.17,0.31) |
| LYCOANNO | 0.52 (0.3,0.83) | 0.3 (0.13,0.7) | 0.52 (0.31,0.83) | 0.18 (0.07,0.48) |
| MAIABIFO | 0.74 (0.67,0.83) | 0.76 (0.62,0.9) | 0.79 (0.72,0.87) | 0.8 (0.67,0.92) |
| MELAPRAT | 0.25 (0.19,0.33) | 0.11 (0.06,0.25) | 0.26 (0.2,0.34) | 0.13 (0.08,0.3) |
| MELASYLV | 0.2 (0.15,0.26) | 0.1 (0.06,0.17) | 0.21 (0.16,0.28) | 0.1 (0.05,0.16) |
| MELINUTA | 0.65 (0.4,0.95) | 0.48 (0.19,0.92) | 0.71 (0.48,0.96) | 0.53 (0.24,0.93) |
| ORTHSECU | 0.08 (0.05,0.16) | 0.13 (0.07,0.18) | 0.09 (0.05,0.17) | 0.09 (0.06,0.12) |
| OXALACET | 0.85 (0.72,0.97) | 0.72 (0.5,0.94) | 0.9 (0.81,0.98) | 0.77 (0.59,0.95) |
| PICEABI3 | 0.34 (0.27,0.43) | 0.14 (0.09,0.18) | 0.36 (0.29,0.45) | 0.13 (0.09,0.18) |
| PINUSYL3 | 0.16 (0.11,0.23) | 0.22 (0.14,0.38) | 0.12 (0.08,0.16) | 0.18 (0.12,0.27) |
| POPUTRE3 | 0.22 (0.13,0.36) | 0.04 (0.03,0.06) | 0.22 (0.14,0.37) | 0.09 (0.04,0.17) |
| PTERAQUI | 0.99 (0.93,1) | 0.83 (0.43,1) | 1 (0.98,1) | 0.94 (0.7,1) |
| RUBUIDA4 | 0.92 (0.78,1) | 0.68 (0.38,0.97) | 0.97 (0.9,1) | 0.94 (0.81,1) |
| RUBUSAXA | 0.91 (0.8,0.99) | 0.8 (0.55,0.98) | 0.94 (0.85,1) | 0.9 (0.7,1) |
| SOLIVIRG | 0.25 (0.06,0.45) | 0.41 (0.23,0.7) | 0.26 (0.07,0.45) | 0.46 (0.26,0.74) |
| SORBAUC3 | 0.28 (0.21,0.35) | 0.34 (0.22,0.53) | 0.29 (0.23,0.37) | 0.39 (0.27,0.56) |
| TRIEEURO | 0.4 (0.34,0.48) | 0.2 (0.16,0.25) | 0.45 (0.38,0.53) | 0.25 (0.19,0.3) |
| VACCMYRT | 0.99 (0.98,0.99) | 0.99 (0.98,1) | 0.99 (0.98,0.99) | 0.99 (0.97,1) |
| VACCULIG | 1 (0.99,1) | 0.99 (0.97,1) | 1 (0.99,1) | 0.99 (0.94,1) |
| VACCVITI | 0.96 (0.95,0.98) | 0.98 (0.96,0.99) | 0.97 (0.96,0.98) | 0.98 (0.96,0.99) |
| VIOLRIVI | 0.09 (0.06,0.12) | 0.05 (0.02,0.08) | 0.15 (0.1,0.26) | 0.22 (0.13,0.36) |

### Figures

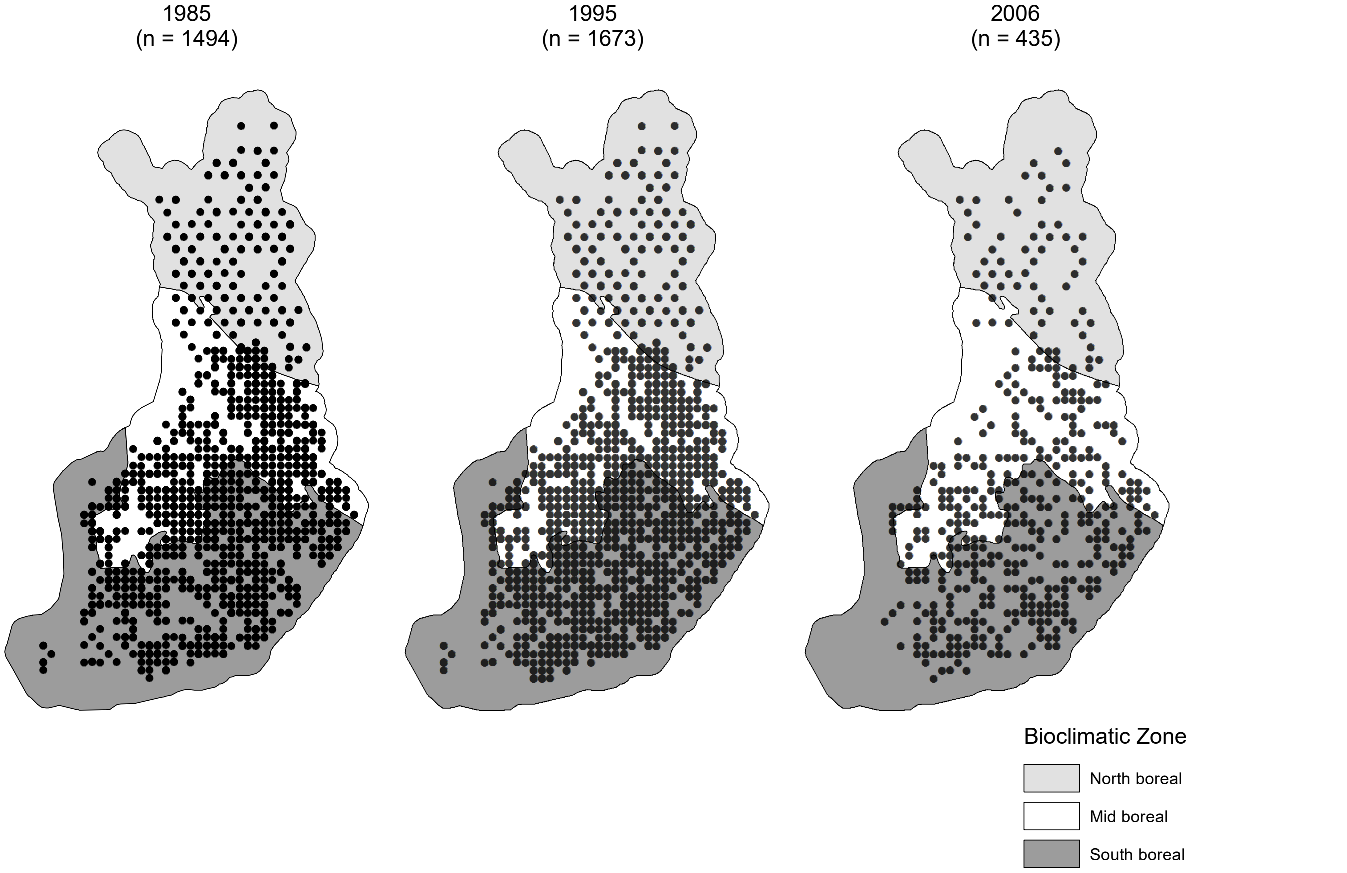


Figure S.1: Understory community inventory sites by year and bioclimatic zone. The overall number of sample sites is provided in parentheses for each year.

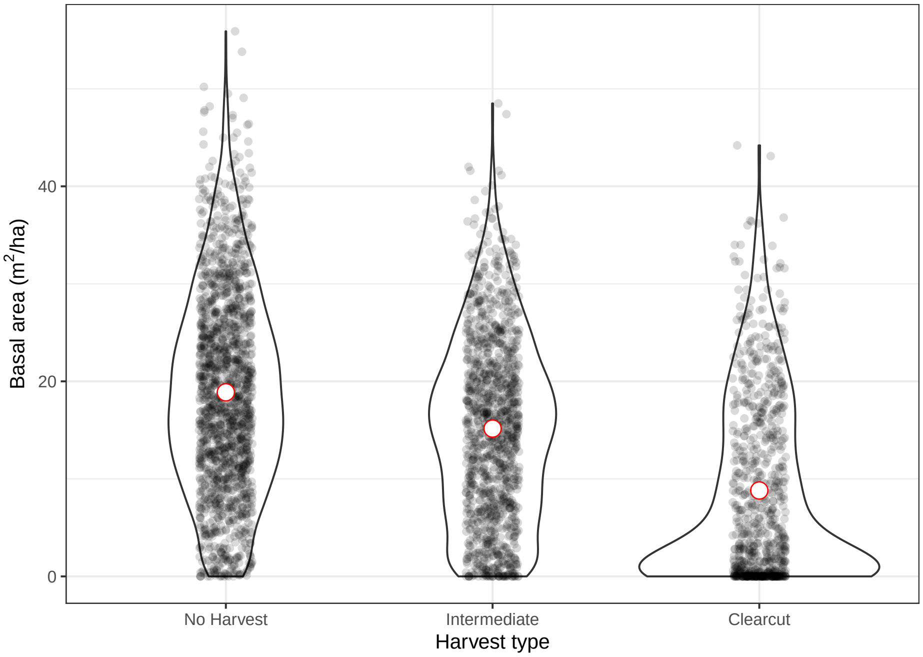


Figure S.2: Basal area per hectare as a function of harvest type within observed understory communities. Jittered site-level observations are overlaid on violin plots of forest density observations. Red points indicate the site-level mean forest density. Intermediate harvest types include forest thinnings and non-clearcut harvests. The clearcut category includes harvests under which all or nearly-all overstory trees are removed. All harvest activity occurred within a 20 year period preceding the inventory year.

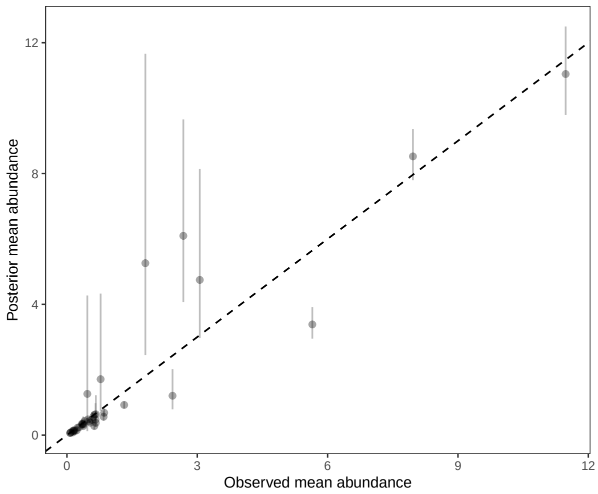


(a) $\mathbf{Y}^{85}|\mathbf{Y}^{95}, \mathbf{Y}^{06}$

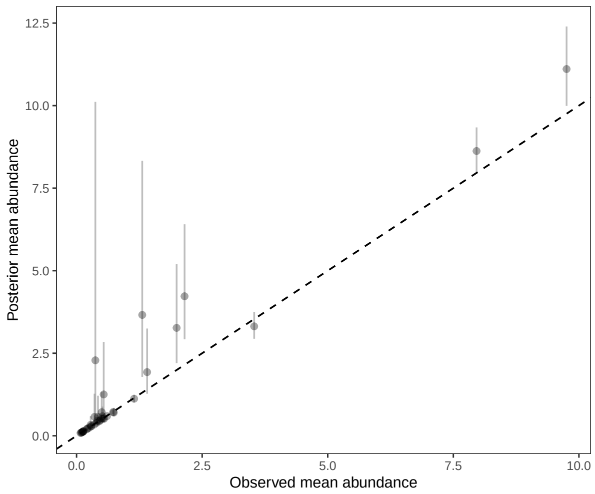


(b) $\mathbf{Y}^{95}, \mathbf{Y}^{06}$

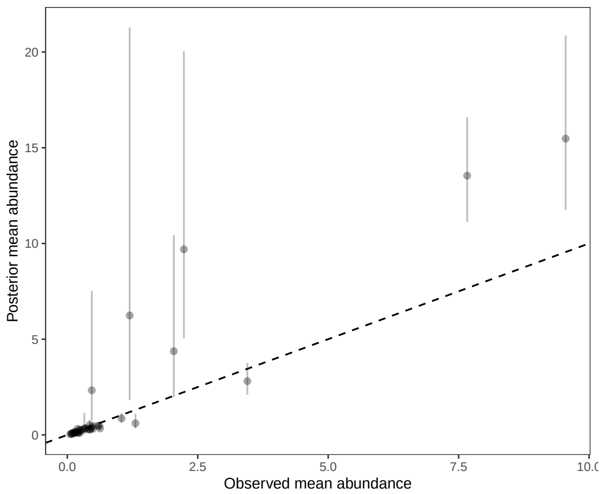


(c) $\mathbf{Y}^{95}|\mathbf{Y}^{06}$

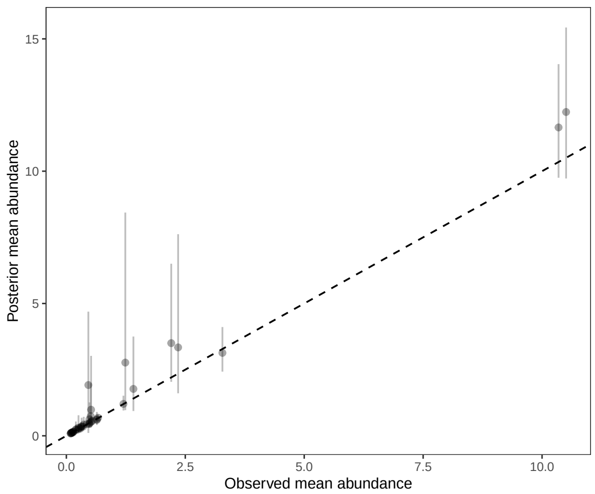


(d) $\mathbf{Y}^{06}$

Figure S.3: Predicted versus observed mean species abundances. Means are calculated across all in-sample or out-of-sample sites. Points correspond to posterior mean values with vertical lines indicating 95 percent credible intervals. A dashed 1-to-1 line is provided for reference. $\mathbf{Y}^{85}|\mathbf{Y}^{95}, \mathbf{Y}^{06}$ and $\mathbf{Y}^{95}|\mathbf{Y}^{06}$ are out-of-sample predictions, while $\mathbf{Y}^{95}, \mathbf{Y}^{06}$ and $\mathbf{Y}^{06}$ are in-sample predictions with the numbers indicating the understory inventory year.
