## Supplementary material for "Bayesian joint species distribution model selection for community-level prediction": Technical appendix

### Additional details on Bayesian JSDM framework

A hierarchical prior is used to estimate species-level responses to the environment, $\boldsymbol{\beta}_j \sim \mathrm{N}(\boldsymbol{\mu}, \mathbf{V})$. Here, $\boldsymbol{\mu}$ defines an among-species mean response to the environment and $\mathbf{V}$ is a covariance matrix (with dimension matching $\mathbf{x}_i$—the number of site-level variables) reflecting covariation in species' responses to the environment (Ovaskainen et al., 2017). The hierarchical prior allows the data to inform the variability of species-level responses to the environment shrinking these to a common mean when there is limited evidence for differential environmental responses (Hooten and Hobbs, 2015).

As noted in the main text, a zero-centered Gaussian distribution is used to model the species random effect term. At a given site $i$, random effects may be modeled independently among species: $w_{ij} \sim \mathrm{N}(0, \sigma_j^2)$ where $\sigma_j^2$ is a species-specific variance term; or dependently among species: $\mathbf{w}_i \sim \mathrm{N}(\mathbf{0}, \boldsymbol{\Omega})$ where $\mathbf{w}_i = (w_{i1}, w_{i2}, \ldots, w_{im})$ and $\boldsymbol{\Omega}$ is an $m$-dimensional, dense covariance matrix often requiring dimension-reduction approaches to be estimated given the number of pairwise species combinations (Ovaskainen et al., 2016, 2017; Taylor-Rodríguez et al., 2017).

### Poisson approximation to multinomial

Under the Poisson approximation to the multinomial (as defined in, McCullagh and Nelder, 1989), the joint abundance of all species at a site is modeled as,

$$\begin{aligned} \mathbf{y}_i &\sim \prod_{j=1}^{m} \mathrm{Pois}(\mu_{ij}) \\ \mu_{ij} &= \exp\left\{\phi_i + \mathbf{x}_i'\boldsymbol{\beta}_j + w_{ij}\right\} \end{aligned} \tag{S.1}$$

where $\phi_i$ is an additive site-specific constant used to ensure that modeled species' abundances sum to the unobserved multinomial sample size (see below), $\mathbf{x}_i$ are site-level variables describing the local environment, $\boldsymbol{\beta}_j$ are species-specific responses to the environment, and $w_{ij}$ is an additive species random effect (the $\mathbf{x}_i$, $\boldsymbol{\beta}_j$, and $w_{ij}$ terms are as defined in the main text). The Poisson model defined in Eqtn. (S.1) can be factored into two components:

$$y_{i\cdot} \sim \mathrm{Pois}(N_i) \tag{S.2a}$$

$$\mathbf{y}_i | y_{i\cdot} \sim \mathrm{Multinom}(y_{i\cdot}, \boldsymbol{\pi}_i). \tag{S.2b}$$

Here, $N_i$ is the mean total abundance of all species defined as $N_i = \sum_{j=1}^{m} \mu_{ij}$, while $\boldsymbol{\pi}_i$ defines the multinomial probability for each species with $\pi_{ij} = \frac{\mu_{ij}}{\sum_{j=1}^{m} \mu_{ij}} = \frac{\exp\left\{\mathbf{x}_i'\boldsymbol{\beta}_j + w_{ij}\right\}}{\sum_{j=1}^{m} \exp\left\{\mathbf{x}_i'\boldsymbol{\beta}_j + w_{ij}\right\}}$. All information for the $\boldsymbol{\beta}_j$'s and $w_{ij}$'s, which fully define the multinomial probabilities, is contained in $\mathbf{y}_i|y_{i\cdot}$ which follows a multinomial distribution. The second component (Eqtn. S.2b) is the model we would like to fit, but the Poisson model defined in Eqtn. (S.1) is the model we actually fit. However, the likelihood functions for the $\boldsymbol{\beta}_j$'s and $w_{ij}$'s are the same under both models, thereby providing equivalent inference on species' probabilities (McCullagh and Nelder, 1989). In a true multinomial data model, the total abundance ($y_{i\cdot}$) is fixed and observed along with the respective abundance of each species whereas under the Poisson approximation, $y_{i\cdot}$ is treated as a random variable with the observed total abundance equivalent to an observation from the underlying Poisson distribution (Eqtn. S.2a). Treating the total abundance as a random variable is sensible in the current analysis given the structure of the understory community data. While we observe species' relative abundances in terms of percent cover, these percent cover values may exceed 100 percent, and a given site may support greater total cover than observed. The sum of all species' abundances represents an observation of the total abundance, but we expect this to be variable and subject to observer error.

### Implementation of Poisson approximation in HMSC

The Poisson approximation to the multinomial (Eqtn. S.1) is implemented in the Hierarchical Model of Species Communities (HMSC) `R` package (Tikhonov et al., 2021) by incorporating a site-specific intercept that is constant with respect to species ($\phi_i$). Details on the HMSC model structure and the Markov chain Monte Carlo (MCMC) sampler used to fit the model are provided in Tikhonov et al. (2020). We assigned a normal prior to the site-specific intercept: $\phi_i \sim \mathrm{N}(\mu_\phi, \tau_\phi^2)$. This prior combined with the likelihood function for the latent linear predictor ($z_{ij} \sim \mathrm{N}(\phi_i + \mathbf{x}_i'\boldsymbol{\beta}_j, \sigma_j^2)$ given $w_{ij} \sim \mathrm{N}(0, \sigma_j^2)$) leads to the following conditional posterior for the site-specific intercept:

$$\begin{aligned}
\phi_i|\mathbf{Y}, \boldsymbol{\theta} &\sim \mathrm{N}(Vv, V) \\
V &= \left( K_i \sum_{j=1}^{m} \frac{1}{\sigma_j^2} + \frac{1}{\tau_\phi^2} \right)^{-1} \\
v &= \sum_{k=1}^{K_i} \sum_{j=1}^{m} \frac{1}{\sigma_j^2} \left( z_{ij}^{(k)} - \mathbf{x}_i'^{(k)} \boldsymbol{\beta}_j \right) + \frac{\mu_\phi}{\tau_\phi^2}
\end{aligned}$$

where $K_i$ is the number of times site $i$ was inventoried based on the response matrix $\mathbf{Y}$ and $\boldsymbol{\theta}$ is used to denote all other model parameters. Under the HMSC notation, $\mathbf{x}_i'\boldsymbol{\beta}_j \equiv L_{ij}^F$ and $w_{ij} \equiv L_{ij}^R$. In most cases, $K_i$ is equal to one, but when fitting models applying both the 1995 and 2006 data, 435 sites are measured twice (once in 1995 and once in 2006). Note that under this approach, we assume the latent total abundance at a site (i.e., the maximum available growing space or community capacity) does not change over time although we observe different values of the total at each inventory. The site-specific intercepts in the Poisson approximation to the multinomial are known to be challenging to estimate (McCullagh and Nelder, 1989). We utilized an informed prior for the intercept term ($\phi_i$) setting $\mu_\phi = 0$ and $\tau_\phi^2 = 0.05$ to ensure it is well identified. These informed hyper-parameters are based on our expectation that on the exponential scale ($\exp \phi_i$), the site-specific intercept will be close to 1.0 corresponding to a total abundance equal to the sum of expected species' abundances. We found the applied hyper-parameters to be sufficiently flexible allowing $\phi_i$ to take on values from 0.25 to over 3.0 while still providing strong convergence of the conditional posterior.

All other model parameters are sampled according to the update steps defined in Tikhonov et al. (2020) with the small adjustment of incorporating the site-specific intercept into all occurrences of the latent linear predictor term ($\mathbf{z}_i$). Specifically, $L_{ij}^F \rightarrow \phi_i + \mathbf{x}_i' \boldsymbol{\beta}_j$. All HMSC model fits applied three MCMC chains each run for 100,000 iterations with a burn-in period of 90,000 samples and a thinning interval of 10 samples such that 1,000 posterior samples were collected from each chain for a total of 3,000 posterior samples. Convergence was assessed visually for all models.

### Log score calculation

In general terms, the log score is expressed as $\log[\mathbf{Y}_{\text{oos}}|\mathbf{Y}_{\text{obs}}] = \log \int_\Theta [\mathbf{Y}_{\text{oos}}|\boldsymbol{\theta}][\boldsymbol{\theta}|\mathbf{Y}_{\text{obs}}]d\boldsymbol{\theta}$ where $\mathbf{Y}_{\text{oos}}$ is a matrix of out-of-sample (oos) natural communities (with dimension $n_{\text{oos}} \times m$), $\mathbf{Y}_{\text{obs}}$ is a matrix of observed (obs) natural communities (with dimension $n_{\text{obs}} \times m$), $\boldsymbol{\theta}$ represents all estimated model parameters, and $[\cdot]$ indicates a probability distribution (Hooten and Hobbs, 2015). We make predictions of out-of-sample communities (a proxy for new data) conditional on our fitted model (as defined by the posterior distribution $[\boldsymbol{\theta}|\mathbf{Y}_{\text{obs}}]$) and integrate over uncertainty in our model parameters reflected in the posterior distribution.

In our study, we used the three understory inventories described in the Natural community data section of the main text to construct two holdout datasets (1995, 1985), and fit each alternative model twice, once using the 2006 inventory data alone to predict 435 communities within the 1995 data ($1995|2006$), and once using the 1995 and 2006 inventory data combined to predict 1,483 communities within the 1985 data ($1985|1995, 2006$). Only a subset of sites within a given inventory year were used for out-of-sample prediction because predictions are available only at sites included in the training data due to the presence of the additive site-specific constant ($\phi_i$) under the Poisson approximation to multinomial (note, site refers to the location of a plot, not the specific community observed at a given time). We estimated the joint log score for the 1985 and 1995 holdout sets (including 1,918 out-of-sample natural communities) as,

$$\ell(\mathbf{Y}_{\text{oos}}^{85}|\mathbf{Y}_{\text{obs}}^{95}, \mathbf{Y}_{\text{obs}}^{06})\ell(\mathbf{Y}_{\text{oos}}^{95}|\mathbf{Y}_{\text{obs}}^{06}) = \sum_{i=1}^{n_{\text{oos}}^{85}} \log[\mathbf{y}_{i,\text{oos}}^{85}|\mathbf{Y}_{\text{obs}}^{95}, \mathbf{Y}_{\text{obs}}^{06}, \mathcal{M}_k] + \sum_{i=1}^{n_{\text{oos}}^{95}} \log[\mathbf{y}_{i,\text{oos}}^{95}|\mathbf{Y}_{\text{obs}}^{06}, \mathcal{M}_k], \tag{S.3}$$

where $\mathbf{y}_{i,\text{oos}}^{\text{year}}$ and $\mathbf{Y}_{\text{obs}}^{\text{year}}$ indicate out-of-sample and observed inventory data from the specified year, $n_{\text{oos}}^{\text{year}}$ indicates the number of out-of-sample sites in the specified year, and $\mathcal{M}_k$ indicates the alternative model used to make predictions (see Table 1 in the main text).

We utilized a sequential predictive (prequential) approach to calculate the joint log score for 1985 and 1995 holdout data conditional on 1995 and 2006 data (Eqtn. S.3). The prequential approach has been shown to provide statistically consistent identification of the true model when applied in conjunction with a proper scoring rule such as the log score (i.e., the probability of selecting the true model converges to 1.0; Dawid and Musio, 2015). The unique structure of the understory inventory data provides a natural partitioning scheme to apply a prequential approach (as outlined above). As such, we preferred to estimate the joint log score using sequential prediction rather than applying cross-validation. We chose to condition the joint log score on the 1995 and 2006 inventory given that it utilizes the most-recent observations and maximizes the number of sites in our out-of-sample data.

The log score in Eqtn. S.3 is "joint" in the sense that it reflects dependence among species within a local community (defined for each row in the $\mathbf{Y}$ matrix and equivalent to the community log score in the main text). This "point-wise by site" approach is only one way we can estimate the joint log score in the context of natural community data where we often have data that is structured among species and locations. These are defined below using general notation.

$$\text{Fully-dependent: } \log[\mathbf{Y}^{\text{oos}}|\mathbf{Y}^{\text{obs}}]$$

$$\text{Point-wise by site: } \sum_{i=1}^{n_{\text{oos}}} \log[\mathbf{y}_i^{\text{oos}}|\mathbf{Y}^{\text{obs}}]$$

$$\text{Point-wise by species: } \sum_{j=1}^{m} \log[\mathbf{y}_j^{\text{oos}}|\mathbf{Y}^{\text{obs}}]$$

$$\text{Fully point-wise: } \sum_{i=1}^{n_{\text{oos}}} \sum_{j=1}^{m} \log[y_{ij}^{\text{oos}}|\mathbf{Y}^{\text{obs}}]$$

The joint log score value varies depending on the specific approach used to calculate it. The fully point-wise approach is most commonly used to estimate the log score (Gelman et al., 2014). In a natural community data setting, this is equivalent to treating species and sites as independent of one another. Note, this is the standard approach used when evaluating JSDMs using cross-validation and common scoring statistics such as the root mean squared prediction error, deviance information criterion, or Watanabe-Akaike information criterion (Norberg et al., 2019). In settings where there is dependence among species and interest is in community-level prediction, the fully-dependent or point-wise by site approaches are preferred since they align with the model objective. Despite its computational advantages, applying the fully point-wise approach when interest is in community prediction may lead to poor conclusions about the performance of alternative models (see also Kettunen et al., 2021).

Given our interest in predicting local (site-level), community composition, we apply the point-wise by site to calculate the joint log score (i.e., the community log score). We treat local communities as independent given species are not considered to be dispersal limited and local communities are sufficiently spaced that we do not have reason to believe that communities closer in space will be more similar than those farther apart. The dependence among species is explicit under models that apply a multinomial data model: $\mathbf{y}_i^{\text{oos}}|y_{i\cdot}, \mathbf{z}_i \sim \text{Multinomial}(y_{i\cdot}, \boldsymbol{\pi}_i)$ with $\pi_{ij} = \frac{\exp z_{ij}}{\sum_{j=1}^m \exp z_{ij}}$. When we assume species are independent of one another, the joint distribution of an out-of-sample community is expressed as the product of independent Poisson distributions, $\mathbf{y}_i^{\text{oos}}|\mathbf{z}_i \sim \prod_{j=1}^m \text{Pois}(\exp z_{ij})$.

### Approximating joint log scores

We applied Monte Carlo integration to estimate the joint log score defined in Eqtn. S.3. Under a multinomial data model, the joint log score is approximated as,

$$\begin{aligned}
&\ell(\mathbf{Y}^{85}_{\text{oos}}|\mathbf{Y}^{95}_{\text{obs}}, \mathbf{Y}^{06}_{\text{obs}})\ell(\mathbf{Y}^{95}_{\text{oos}}|\mathbf{Y}^{06}_{\text{obs}}) \approx \\
&\sum_{i=1}^{n^{85}_{\text{oos}}} \log\left\{\frac{1}{S}\sum_{s=1}^{S} \text{Multinom}\left(\mathbf{y}^{85}_{i,\text{oos}}|\boldsymbol{\pi}_i^{85|95,06(s)}, y^{85}_{i\cdot,\text{oos}}\right) \text{Pois}\left(y^{85}_{i\cdot,\text{oos}}|N_i^{85|95,06(s)}\right)\right\} \\
&+\sum_{i=1}^{n^{95}_{\text{oos}}} \log\left\{\frac{1}{S}\sum_{s=1}^{S} \text{Multinom}\left(\mathbf{y}^{95}_{i,\text{oos}}|\boldsymbol{\pi}_i^{95|06(s)}, y^{95}_{i\cdot,\text{oos}}\right) \text{Pois}\left(y^{95}_{i\cdot,\text{oos}}|N_i^{95|06(s)}\right)\right\}.
\end{aligned} \tag{S.4}$$

Here, $n^{\text{year}}_{\text{oos}}$ indicates the total number of out-of-sample inventory sites in the specified inventory year (1995, 1985), $\mathbf{y}^{\text{year}}_{i,\text{oos}}$ and $y^{\text{year}}_{i\cdot,\text{oos}}$ indicate the relative species abundances and total abundance, respectively, for the out-of-sample natural community at the $i$th site in the listed inventory year, the $v_i^{\text{year}_{\text{oos}}|\text{year}_{\text{obs}}(s)}$ notation indicates the $s$th posterior sample of $v_i^{\text{year}_{\text{oos}}}$ from the model fit using observed data for the specified inventory year(s), $\boldsymbol{\pi}_i$ represents the multinomial probabilities as defined in Eqtn. (2) of the main manuscript, $N_i = \sum_{j=1}^m \exp\left\{\phi_i + \mathbf{x}_i'\boldsymbol{\beta}_j + w_{ij}\right\}$ as defined in Eqtn. (S.1), and $S$ is the total number of posterior samples (the same number of samples is used for each model fit). Note that the log score calculated under Eqtn. (S.4) is for the joint distribution of $\mathbf{y}_i$ and $y_{i\cdot}$, the relative species composition and total abundance at a site, respectively.

Under the conditionally independent Poisson model, the joint log score is approximated as,

$$\begin{aligned}
\ell(\mathbf{Y}^{85}_{\text{oos}}|\mathbf{Y}^{95}_{\text{obs}}, \mathbf{Y}^{06}_{\text{obs}})\ell(\mathbf{Y}^{95}_{\text{oos}}|\mathbf{Y}^{06}_{\text{obs}}) \approx &\sum_{i=1}^{n^{85}_{\text{oos}}} \log\left\{\frac{1}{S}\sum_{s=1}^{S}\left\{\prod_{j=1}^{m} \text{Pois}\left(y^{85}_{ij,\text{oos}}|z_{ij}^{85|95,06(s)}\right)\right\}\right\} \\
&+\sum_{i=1}^{n^{95}_{\text{oos}}} \log\left\{\frac{1}{S}\sum_{s=1}^{S}\left\{\prod_{j=1}^{m} \text{Pois}\left(y^{95}_{ij,\text{oos}}|z_{ij}^{95|06(s)}\right)\right\}\right\}.
\end{aligned} \tag{S.5}$$

Here, $z_{ij}^{\text{year}_{\text{oos}}|\text{year}_{\text{obs}}(s)}$ indicates the $s$th posterior sample of the latent linear predictor $z_{ij}^{\text{year}_{\text{oos}}}$ from the model fit using observed data for the specified inventory year(s) and $j$ indexes the species ($j = 1, \ldots, m$).

Given concern over the stability of Monte Carlo integration under our high-dimensional, structured data setting, we simulated confidence intervals for the log score using a non-parametric bootstrap of the posterior samples. Specifically, we estimated the joint log score applying Eqtns. (S.4, S.5), depending on the data model, 1,000 times re-sampling 500 posterior samples with replacement from the total number of posterior samples collected.

### Evaluating predictions within data partitions

We assessed the ability of four models (stochastic-independent, environment-independent, stochastic-compositional, environment-compositional; see Table 1 in main text) to predict natural communities and the abundances of individual species within each of our eight defined data partitions (see Model selection section). We again utilized the log score to quantify predictive performance, but calculated the mean log score across sites within each data partition rather than the combined log score to measure the average ability of evaluated models.

The mean, per-site log score for community-level prediction is given by:

$$\bar{\ell}(\mathbf{y}(q)|\mathbf{Y}^{06}_{\text{obs}}, \mathbf{Y}^{95}_{\text{obs}}) = \left(\frac{1}{n^{85}_{q,\text{oos}} + n^{95}_{q,\text{oos}}}\right) \left\{ \sum_{i=1}^{n^{85}_{q,\text{oos}}} \log[\mathbf{y}^{85}_{i,\text{oos}}(q)|\mathbf{Y}^{95}_{\text{obs}}, \mathbf{Y}^{06}_{\text{obs}}, \mathcal{M}_k] + \sum_{i=1}^{n^{95}_{q,\text{oos}}} \log[\mathbf{y}^{95}_{i,\text{oos}}(q)|\mathbf{Y}^{06}_{\text{obs}}, \mathcal{M}_k] \right\} \tag{S.6}$$

where $q$ indexes the data partition such that $\mathbf{y}(q)$ refers to a natural community in the $q$th partition.

The mean, per-site log score for species-level prediction is given by:

$$\bar{\ell}(y_j(q)|\mathbf{Y}^{06}_{\text{obs}}, \mathbf{Y}^{95}_{\text{obs}}) = \left(\frac{1}{n^{85}_{q,\text{oos}} + n^{95}_{q,\text{oos}}}\right) \left\{ \sum_{i=1}^{n^{85}_{q,\text{oos}}} \log[y^{85}_{ij,\text{oos}}(q)|\mathbf{Y}^{95}_{\text{obs}}, \mathbf{Y}^{06}_{\text{obs}}, \mathcal{M}_k] + \sum_{i=1}^{n^{95}_{q,\text{oos}}} \log[y^{95}_{ij,\text{oos}}(q)|\mathbf{Y}^{06}_{\text{obs}}, \mathcal{M}_k] \right\} \tag{S.7}$$

where $q$ again indexes the data partition such that $y_j(q)$ refers to the abundance of a constituent species ($j = 1, \ldots, m$) within the $q$th partition. To estimate species-level log scores under multinomial community models, we apply a Poisson data model for each species (the marginal species distribution under the Poisson approximation to multinomial): $y_{ij} \sim \text{Pois}\left(\exp\left\{\phi_i + \mathbf{x}_i'\boldsymbol{\beta}_j + w_{ij}\right\}\right)$.

In practice, we apply Monte Carlo integration to approximate the mean log scores for natural communities and constituent species in each data partition updating Eqtn. (S.4) to reflect the out-of-sample data in the $q$th partition and the target of prediction (community or constituent species).

### Model checking

To evaluate model fit, Bayesian R-squared values were calculated for each species under the best performing environment-compositional model applying the methodology presented in Gelman et al. (2019). Specifically, for each posterior sample we calculated the among-site variance of the expected abundance of each species ($V_j^{\text{mod}(s)}$) as well as the mean residual variance at each site ($V_j^{\text{res}(s)}$) where $(s)$ indexes the posterior sample. Under the environment-compositional model applying the Poisson approximation to multinomial, the variance of the expected abundance of a species was calculated as, $V_j^{\text{mod}(s)} = \frac{1}{n_k-1}\sum_{i=1}^{n_k}\left(N_i^{(s)}\pi_{ij}^{(s)} - \frac{1}{n_k}\sum_{i=1}^{n_k} N_i^{(s)}\pi_{ij}^{(s)}\right)^2$, while the mean residual variance for a species at a given site was calculated as, $V_j^{\text{res}(s)} = \frac{1}{n_k}\sum_{i=1}^{n_k} N_i^{(s)}\pi_{ij}^{(s)}(1-\pi_{ij}^{(s)})$ where $n_k$ is the number of sample sites in a given understory inventory. The Bayesian R-squared value for the $s$th posterior sample was the calculated as: $r^{2(s)} = \frac{V_j^{\text{mod}(s)}}{V_j^{\text{mod}(s)}+V_j^{\text{res}(s)}}$ (Gelman et al., 2019). Bayesian R-squared values were calculated for both the in-sample and out-of-sample data of each model fit (Table S.4).

We further generated plots of the posterior predictive distribution of species' abundances under the environment-compositional model relative to observed quantities. We applied composition sampling to sample from the posterior predictive distribution for a local community. Specifically, we sampled $y_{i\cdot}^{(s)} \sim \text{Pois}(N_i^{(s)})$ and $\mathbf{y}_i^{(s)}|y_{i\cdot}^{(s)} \sim \text{Multinom}(y_{i\cdot}^{(s)}, \boldsymbol{\pi}_i^{(s)})$ for each posterior sample $(s = 1, \ldots, S)$. Posterior predictive samples were summarized by species and plotted relative to observed species' abundances for both the in-sample and out-of-sample data of each model fit (Fig. S.3).
